# Type I interferon induces TCR-dependent and -independent antimicrobial responses in γδ intraepithelial lymphocytes

**DOI:** 10.1101/2024.03.11.584444

**Authors:** Matthew A. Fischer, Luo Jia, Karen L. Edelblum

## Abstract

Intraepithelial lymphocytes (IEL) expressing the γδ T cell receptor (TCR) survey the intestinal epithelium to limit the invasion of microbial pathogens. The production of type I interferon (IFN) is a central component of an antiviral immune response, yet how these pro-inflammatory cytokines contribute to γδ IEL effector function remains unclear. Based on the unique activation status of IELs, and their ability to bridge innate and adaptive immunity, we investigated the extent to which type I IFN signaling modulates γδ IEL function. Using an *ex vivo* culture model, we find that type I IFN alone is unable to drive IFNγ production, yet low level TCR activation synergizes with type I IFN to induce IFNγ production in murine γδ IELs. Further investigation into the underlying molecular mechanisms of co-stimulation revealed that TCRγδ-mediated activation of NFAT and JNK is required for type I IFN to promote IFNγ expression in a STAT4- dependent manner. Whereas type I IFN rapidly upregulates antiviral gene expression independent of a basal TCRγδ signal, neither tonic TCR triggering nor the presence of a TCR agonist was sufficient to elicit type I IFN-induced IFNγ production *in vivo*. However, bypassing proximal TCR signaling events synergized with IFNAR/STAT4 activation to induce γδ IEL IFNγ production. These findings indicate that γδ IELs contribute to host defense in response to type I IFN by mounting a rapid antimicrobial response independent of TCRγδ signaling, and under permissive conditions, produce IFNγ in a TCR-dependent manner.

## Introduction

The intestinal barrier is composed of a single layer of epithelial cells that separates the underlying immune compartment from luminal microbes. Disruption of this barrier can lead to microbial invasion and activation of mucosal immunity; therefore, maintenance of epithelial integrity and immunosurveillance is critical to prevent an inflammatory response (1–3). Tissue-resident intraepithelial lymphocytes (IEL) expressing the γδ TCR patrol the epithelium to facilitate a rapid response to pathogens (4–6). Among the most well-characterized innate immune responses to microbes is the production of type I IFN (7–9), a family of pro-inflammatory cytokines that signal through the IFNα/β receptor (IFNAR) leading to activation and phosphorylation of STAT1/2 heterodimers that complex with interferon regulatory factor (IRF) 9, or phosphorylation of STAT4 homodimers (10–12). In contrast, type II IFN, or IFNγ, signals through a different receptor with unique transcriptional targets. In the context of infection, IFNγ secretion by NK cells acts as a link between innate immunity and the activation of an adaptive response (13). Previous studies show that γδ IELs produce type I, II, and III IFNs in response to stimulation of the TCR complex with agonist antibodies *in vitro*, and that TCR activation *in vivo* protects mice against viral infection in a type I/III IFN-dependent manner (14). Although TCR activation induces IFN production, the extent to which type I IFN-mediated signaling regulates

γδ IEL effector responses remains unclear.

IELs can be classified into two main subsets: induced IELs, which arise from conventional CD8αβ or CD4 T cells and home to the intestinal epithelium in response to cognate antigen, and natural IELs that express CD8αα (either TCRαβ or TCRγδ) and can be activated in an MHC-independent manner (15–17). Under homeostatic conditions, γδ IELs remain poised in an activated, yet immunologically quiescent state (18–20), reflecting prior observations that the TCR is constantly triggered *in vivo* (21, 22). Based on the rapid and conserved response of γδ IELs to microbial invasion (4, 14), we aimed to elucidate the underlying mechanisms by which type I IFN may influence γδ IEL effector function.

Type I IFN has been widely studied in the context of viral infection, in which the timing and stoichiometry of STAT1 and STAT4 expression, modulate the innate and adaptive arms of the host response. During early infection, high STAT1 levels prevent non-specific CD8 T cells from proliferating and producing IFNγ (23–26). Once activated, the upregulation of STAT4 contributes to CD8 T cell expansion and pro-inflammatory cytokine production to promote viral clearance. In contrast, high levels of STAT4 at steady-state allows NK cells to rapidly respond to infection; however, these pro-inflammatory responses are restricted during the adaptive phase of infection through STAT1-mediated mechanisms (27–30). Based on the unique activation status of IELs and functional similarities between unconventional γδ T cells and NK cells, we investigated the extent to which IFNAR signaling modulates γδ IEL function as these sentinel lymphocytes are likely exposed to both tonic and pathogenic levels of type I IFN in the intestinal mucosa (7).

In this study, we show that type I IFN enhances γδ IEL IFNγ production in a co-stimulatory manner *ex vivo*, whereas treatment with type I IFN alone is unable to elicit cytokine production. Suboptimal concentrations of TCR agonist were sufficient to drive type I IFN- mediated IFNγ production through the activation of NFAT and JNK. Moreover, we find that IFNAR activation of STAT4 synergizes with TCR signaling to enhance IFNγ production.

Transcriptomic profiling of γδ IELs revealed that type I IFN induced the upregulation of antimicrobial gene programs, including interferon-stimulated genes (ISG), independent of TCR signaling. We found that basal TCR triggering was inadequate for γδ IEL IFNγ induction *in vivo*, and moreover, all natural CD8αα IELs exhibited limited IFNγ production *in vivo*. However, bypassing proximal TCR signaling events induced IFNγ in freshly-isolated γδ IELs, and subsequently, resulted in increased IFNγ production following IFNα exposure in a STAT4- dependent manner. Taken together, these findings indicate that γδ IELs contribute to antimicrobial host immunity in response to type I IFN through rapid TCR-independent ISG expression, and under permissive conditions, may also promote TCR-dependent IFNγ production.

## Materials and Methods

Animals. Experiments were performed on 8–12-week-old mice of both sexes on a C57BL/6 background. Nur77-GFP (C57BL/6-Tg Nr4a1-EGFP/cre^820Khog^/J) and "interferon-gamma reporter with endogenous polyA transcript" (GREAT)(B6.129S4-*Ifng^tm3.1Lk^y*/J) mice were obtained from The Jackson Laboratory. TcrdH2BEGFP (TcrdEGFP) mice were provided by Bernard Malissen (INSERM)(31) and crossed with STAT1 KO or STAT4 KO mice provided by Sergei Kotenko or Tessa Bergsbaken, respectively (Rutgers New Jersey Medical School). Mice were housed in a room with monitored temperature and humidity for a 12-hour light/dark cycle in cages with aspen shavings, autoclaved 5010 chow, and tap water. All mice except Nur77-GFP were crossed to TcrdEGFP mice and thus would have been exposed to similar microbiota. For experiments using Nur77-GFP mice, wildtype (WT) littermate controls were used. For TCRγδ inhibition, mice were treated with 200 ÿg anti-TCRγδ (UC7-13D5) or Armenian Hamster IgG (BioXCell, BE0091) antibody i.p. 48 and 24 hours prior to the experiment. Mice were treated with 1 ÿg murine IFNα2 (Bon Opus Biosciences) or endotoxin-free PBS (Fisher Scientific) via retroorbital injection 5 hours prior to euthanasia. Alternatively, mice were treated with 25 ÿg anti-CD3 (145-2C11, Biolegend) or Armenian Hamster IgG and 1 ÿg murine IFNα2 i.p. for 24 hours. All studies were conducted in an Association of the Assessment and Accreditation of Laboratory Animal Care-accredited facility according to protocols approved by Rutgers New Jersey Medical School Comparative Medicine Resources or the Center for Comparative Medicine and Surgery at Icahn School of Medicine at Mount Sinai.

### IEL isolation and *ex vivo* culture

Small intestinal IELs were isolated as described previously (32). Briefly, mice were euthanized after which the small intestine was dissected and opened longitudinally following removal of Peyer’s patches. One cm pieces of small intestine were then incubated in HBSS containing 3 mM EDTA and 7.5% FBS (ThermoFisher) for 1 hour in a shaker at 37° C. Supernatants were collected and IELs enriched using a 20/45/70% Percoll density gradient (Cytiva).

γδ IELs were cultured *ex vivo* as previously described (33) in which 100,000 γδ IELs were activated with 1 ÿg/mL anti-CD3 (2C11, BioLegend) in the presence of 10 U/mL IL-2, 100U/mL IL-3, 200 U/mL IL-4, and 200 ng/mL IL-15 (PeproTech) for 48 h. IELs were then rested in media supplemented with 20 U/mL IL-2 and 100 ng/mL IL-15 for 5 days prior to activation. IELs were re-stimulated with 1 ÿg/mL anti-CD3, 10 ng/mL IFNA/D (PBL Assay Science), 10 ng/mL IL-12 (PeproTech), or a combination of anti-CD3 and cytokine for 5 h. Cultures were supplemented with Golgi Plug (BD Biosciences) for the last 3 h. Alternatively, sorted γδ IELs were treated with 4 ÿg/mL ionomycin (Sigma-Aldrich), 40 ng/mL PMA (Sigma-Aldrich), or both in the presence or absence of 10 ng/mL IFNA/D for 5 h and supplemented with Golgi Plug for the final 4 h. For experiments using pharmacological inhibitors, γδ IELs were pre-treated with each inhibitor for 30 minutes at 37° C prior to stimulation. Inhibitors were used at the following concentrations: 10 ÿM SP600125, 1 ÿM VTX27, 10 ÿM U0126, 10 ÿM LY294002, 10 ÿM INCA6, or 5 ÿM Cyclosporin A.

### Flow cytometric analysis

Flow cytometric data was collected using a BD LSRFortessa X-20 and analyzed using FlowJo software (FlowJo, v10.9.0). IELs were stained in PBS supplemented with Brilliant Stain Buffer (BD Biosciences) as needed. For intracellular staining, cells were fixed and permeabilized with BD Cytofix/Cytoperm solution. For phospho-staining, IELs previously stained with extracellular markers were fixed and permeabilized in eBioscience Foxp3/Transcription Factor Fixation/Permeabilization buffer followed by 30 min nuclear permeabilization with 100% methanol on ice. Intracellular staining was performed using eBioscience Foxp3/Transcription Factor Staining buffer. When necessary, cells were sorted based on GFP or TCRγδ expression using a BD FACSAria Fusion sorter with 95-98% purity. The following antibodies for flow cytometry were used from Biolegend: CD4 (RM4-5), CD44 (IM7), CD69 (H1.2F3), CD8b (YTS156.7.7), CD107a (1D4B), Tcrb (H57-597), Tcrd (GL3), IFNγ (XMG1.2), or BD Biosciences: CD3ε (145-2C11), CD8α (53-6.7), CD103 (M290), STAT1 Y701 (4a), STAT4 Y693 (38/p-Stat4), TNF (MP6-XT22). eBioscience Fixable Viability Dye eFluor 780 was used as a live-dead marker.

### RNA extraction, library preparation, and sequence analysis

RNA was isolated from 5x10^5^ GFP γδ IELs, extracted using TRIzol (Invitrogen) and purified using RNeasy Mini Kit (Qiagen) following chloroform extraction. Following reverse transcription using an iScript cDNA Synthesis Kit (Bio-Rad), qPCR was performed using SYBR Green (Fisher Scientific) on the QuantStudio 6 platform (ThermoFisher Scientific). Ct values of *Ifng* were normalized to the Ct values for *Gapdh*. Primers: *Ifng* (F- GATGCATTCATGAGTATTGCCAAGT; R- GTGGACCACTCGGATGAGCTC), *Gapdh* (F- AAGGTGGTGAAGCAGGCATCTGAG; R- GGAAGAGTGGGAGTTGCTGTTGAAGTC) For bulk RNAseq, γδ IELs were isolated, sorted, and RNA was isolated as described above. SMART-Seq HT kit (Takara) was used for full-length cDNA synthesis and amplification followed by library preparation using Illumina Nextera XT kit. Sequence libraries were multiplexed and sequencing was performed using a 2x 150 paired end configuration (Illumina HiSeq). Raw data files were converted to fastq files, demultiplexed using bcl2fastq 2.20 software, and analyzed using Partek Flow pipeline (Partek, PGS7.21.1119). In Partek, reads were aligned to the most recently annotated *Mus musculus* (mm39) genome using STAR aligner (version 2.7.8a). A mean of 34,538,531 read pairs (87.9%) were uniquely mapped to the mm39 genome. Aligned reads were assigned to genomic features using the Partek E/M annotation model and annotated with Ensembl transcript (release 104). Genes with a maximum read count ≤100 reads were filtered out and remaining reads were normalized to the median ratio. DESeq2 was used to determine differential expression between IgG-IFNα, UC7-PBS, or UC7-IFNα treated and IgG-PBS treated samples. Differential gene expression (DEG) was filtered based on a false discovery rate (FDR) < 0.05 and a fold change of ± 1.5. Upregulated DEGs were compared to a list of upregulated ISGs from published microarray data of splenic γδ T cells following 2 h treatment with 10,000 U IFNα *in vivo* (34). Gene set enrichment (GSE) used the 2023-09-13 Gene Ontology database (GO Consortium) to identify GO terms associated with up or downregulated DEGs with a *p*<0.05.

### Statistical analyses

Data is represented as the mean ± SEM. Statistical analysis was performed using Prism (GraphPad, v10). Unpaired t-tests were used to compare between two conditions, whereas one-way or two-way ANOVA with either Tukey’s or Dunnett’s post hoc test were used to compared between more than two conditions. A single experimental replicate with 3-4 mice per group was performed for the transcriptomics studies and all other experiments were performed with least two experimental replicates. For *ex vivo* studies, technical replicates were performed with primary cells isolated from a single mouse split between all experimental conditions, and thus each data point represents an individual mouse.

## Results

### Type I IFN enhances γδ IEL IFNγ production

To investigate the effects of IFNα on γδ IEL effector cytokine production, we took advantage of an *ex vivo* culture model to measure the response to type I IFN in the presence or absence of a TCR agonist antibody. Briefly, γδ IELs were stimulated for 2 days in the presence of anti-apoptotic and pro-proliferative cytokines and rested for 5 days prior to restimulation (14). As expected, γδ IELs rapidly respond to TCR stimulation as measured by an increase in the frequency of IFNγ^+^ cells, as well as the frequency of TNF^+^ cells, CD107a externalization, and TCRγδ internalization compared to untreated cells (Figure 1, Supplementary Figure 1).

**Figure 1:**
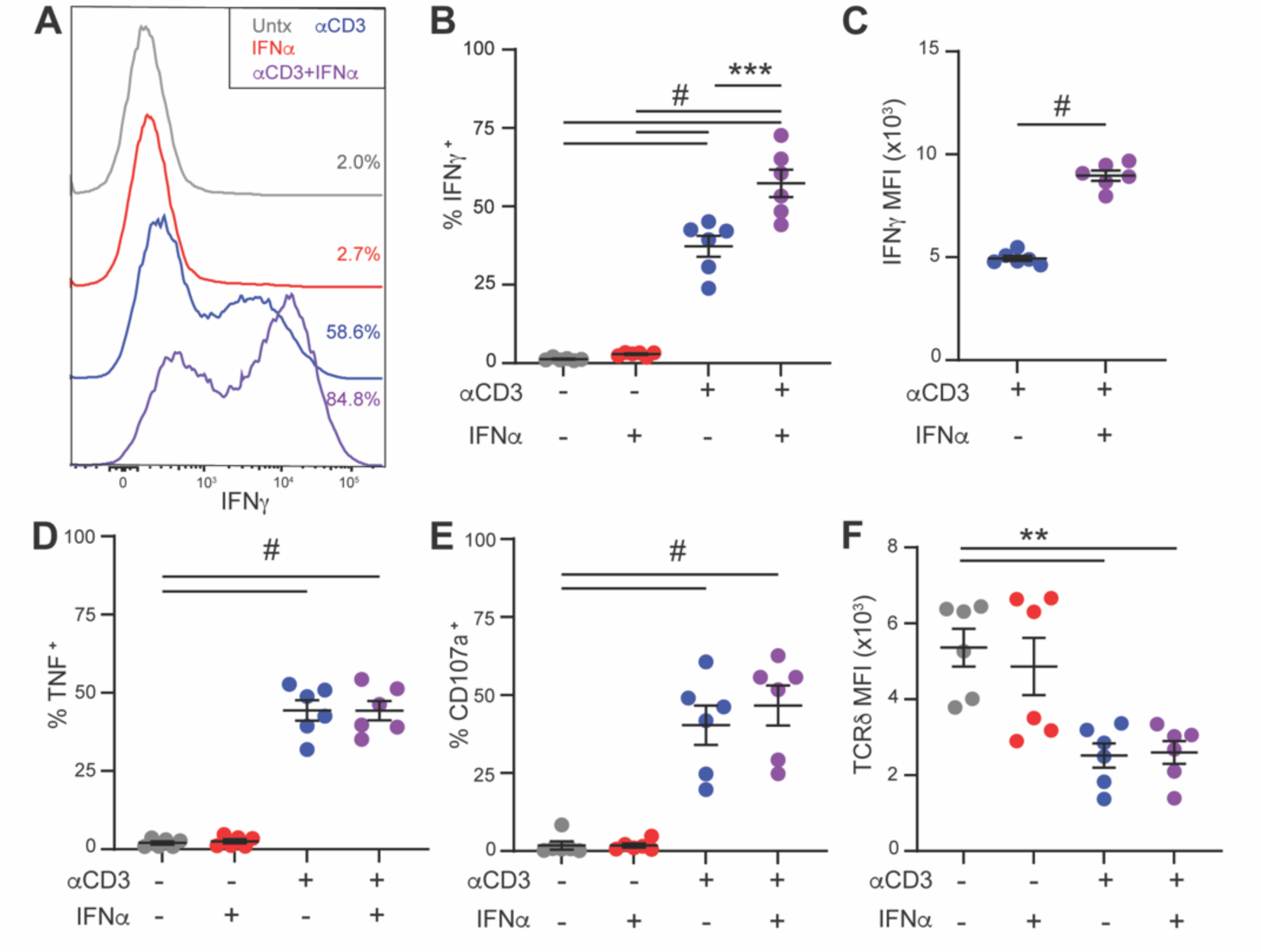
TCR activation is required for type IIFN mediated-enhanced IFNγ production in γδ lELs. Flow cytometry was performed on ex *vivo* cultured γδ lELs treated with 1 µg/mL ctCD3, 10 ng/mL IFNα, or both for 5 h. (A) Representative histogram showing (B) the frequency of IFNγ* γδ lELs and (C) IFNγ MFI. The frequency of (D) TNF^+^ or (E) CD107a^+^ γδ lELs or (D) TCRδ MFI is shown. Data are graphed as mean ± SEM. Two independent experiments; n=6 mice; **p <0.01, ***p <0.001, #p <0.0001; B, D-F: One-way ANOVA with Tukey’s multiple comparison test, C: Unpaired t test.

Treatment with IFNα alone did not induce a measurable effector response; however, treatment with IFNα and TCR agonist resulted in enhanced IFNγ compared to anti-CD3 alone (Figure 1A- C). The co-stimulatory effect of IFNα on γδ IELs was unique to IFNγÿproduction, as combined exposure had no effect on TNF production or CD107a externalization relative to TCR stimulation alone (Figure 1D,E). Similar results were observed at the transcript level (Supplementary Figure 2A), demonstrating that IFNα influences the transcriptional regulation of *Ifng* as opposed to modulating protein translation. Taken together, these data indicate that IFNα facilitates a more robust IFNγ transcriptional response by γδ IELs following TCR activation *ex vivo*.

### Suboptimal TCR activation is sufficient to facilitate γδ IEL effector response following IFNα exposure

Since IFNα requires the presence of a TCR signal to induce IFNγ production (Figure 1), we next asked whether type I IFN directly affects the magnitude of TCRγδ signaling. To this end, we isolated γδ IELs from Nur77-GFP reporter mice, which express GFP when Nur77 is activated downstream of the TCR (35). No change in the MFI of Nur77-GFP was observed following TCR activation regardless of IFNα exposure (Figure 2A), demonstrating that type I IFN does not influence TCR signal strength. Further analysis confirmed that only Nur77-GFP^+^ cells produce cytokine and that the co-stimulatory effect of IFNα on IFNγ production is specific to γδ IELs responding to a TCR agonist (Figure 2B,C). Again, IFNα treatment had no effect on TNF production (Figure 2C).

**Figure 2:**
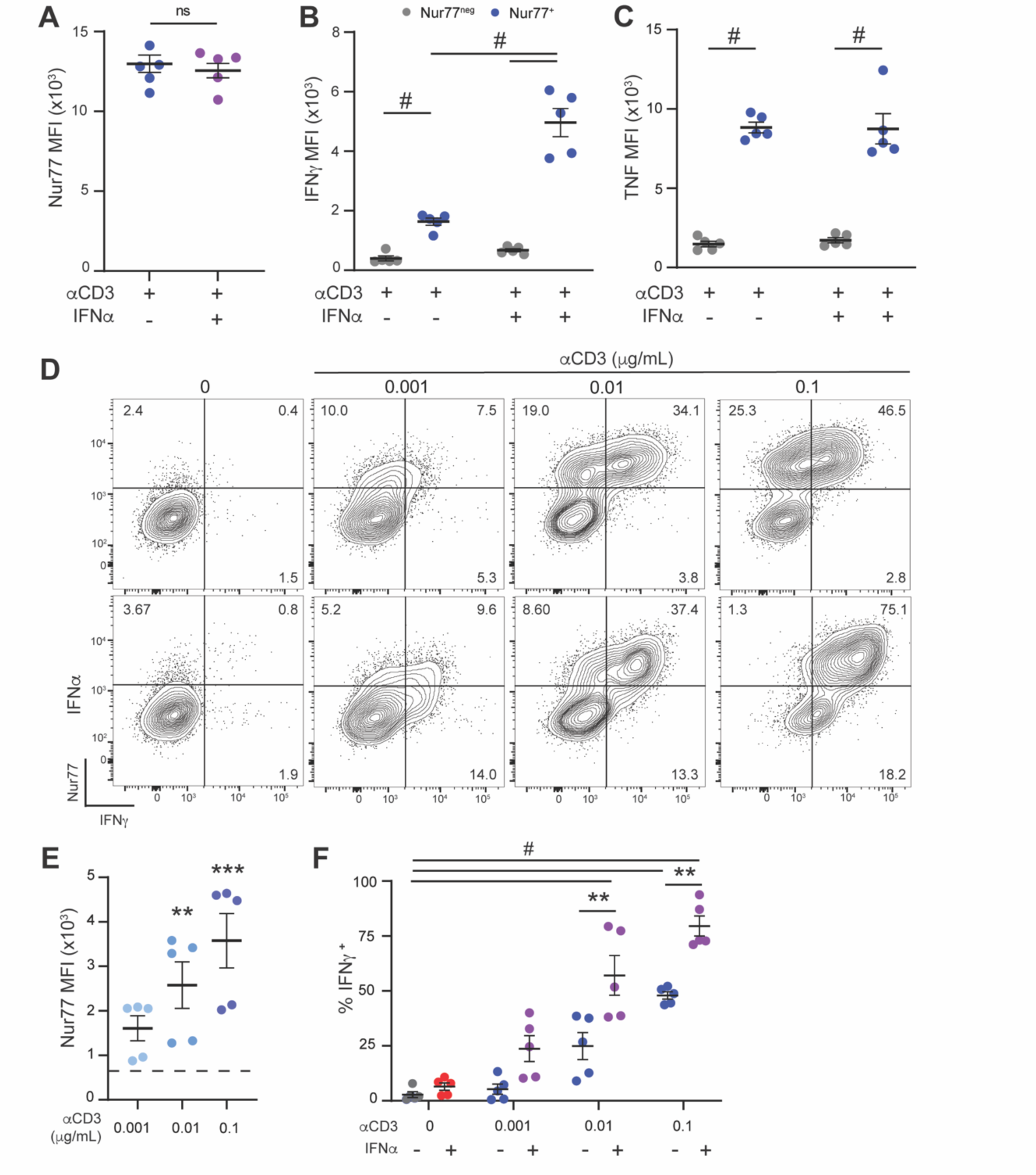
Suboptimal TCRγδ signaling is sufficient for type I IFN-mediated IFNγ production. γδ lELs were sorted from Nur77-GFP and stimulated ex *vivo* with 1µg/mLαCD3, 10 ng/mL IFNα, or both for 5 h. (A) MFI of Nur77-GFP from γδ lELs treated with αCD3 ± IFNα. The MFI of (B) IFNγ or (C) TNF in Nur77* or Nur77^n^« γδ lELs. (D) Representative flow plot of Nur77 and IFNγ expression in γδ lELs treated with 0-0.1 µg/mL αCD3 for 5 h in the presence or absence of 10 ng/mL IFNα. (E) MFI of Nur77 in γδ lELs treated with 0-0.1 µg/mL αCD3 for 5 h . Dashed line represents the average Nur77 MFI among untreated cells (Nur77^nθs^) used for statistical comparison. (F) Frequency of IFNγ* γδ lELs following treatment with 0.001-0.1 µg/mL αCD3 ± IFNα. Data are shown as mean ± SEM. Two independent experiments; n=5-6 mice; **p <0.01, ***p <0.001, #p <0.0001; A: Unpaired t test, B,C,F: Two-way ANOVA with Tukey’s multiple comparison test, E: One-way ANOVA with Tukey’s multiple comparison test.

γδ IELs are immunologically quiescent under homeostatic conditions; therefore, we hypothesized that a certain threshold of TCR signaling must be achieved to observe cytokine production in response to IFNα. While increasing concentrations of TCR agonist elicited a dose-dependent response in Nur77 expression (Figure 2D,E), we found that stimulation with as little as 0.01ÿg/ml anti-CD3 resulted in IFNα-mediated IFNγ production (Figure 2F). Complementing these findings, we found that a low concentration of IFNα (0.1 ng/mL) was sufficient to enhance IFNγ production in response to 0.1 ÿg/mL anti-CD3, the lowest dose of TCR agonist demonstrating a significant increase in IFNγ in both anti-CD3 and co-stimulated conditions relative to control (Supplementary Figure 2B, Figure 2F). Together, these data show that suboptimal concentrations of TCR agonist are sufficient to surpass the γδ IEL activation threshold, thus allowing type I IFN to enhance IFNγ production *ex vivo*.

### Activation of NFAT and JNK is required for γδ IEL IFNγ production in response to TCR and IFNα stimulation

Our data suggests that IFNα enhances the transcription of *Ifng* through a mechanism that is parallel to, and dependent upon, TCR activation. In conventional T cells, TCR ligation and CD28 co-stimulation results in the activation of multiple downstream pathways, including calcium, NF- κB, MAPK, and PI3K signaling, culminating in the phosphorylation and/or translocation of various transcription factors that induce T cell effector programs (36). However, unlike conventional T cells, CD28 is not expressed on γδ IELs, which may contribute to their reduced responsiveness to TCR stimulation (37). Therefore, to elucidate the TCR-mediated signaling pathways contributing to enhanced IFNγ production in response to IFNα, we treated freshly- isolated γδ IELs with IFNα in the presence or absence of PMA and/or ionomycin (P/I) to differentially activate pathways downstream of TCR signaling. Ionomycin facilitates the release of intracellular calcium to activate calcineurin/NFAT, whereas PMA is a diacylglycerol (DAG) analog that induces AP-1 and NF-κB signaling (38). Activation of γδ IELs with ionomycin promoted IFNγ expression in response to IFNα, whereas PMA alone failed to stimulate a response (Figure 3A,B). Notably, treatment with either ionomycin or PMA was unable to stimulate IFNγ production, yet in combination, γδ IELs produced IFNγ that was further enhanced with IFNα co-stimulation. These data suggest that ionomycin-induced calcium release in combination with a secondary PMA-induced signal contributes to γδ IEL IFNγ production.

**Figure 3:**
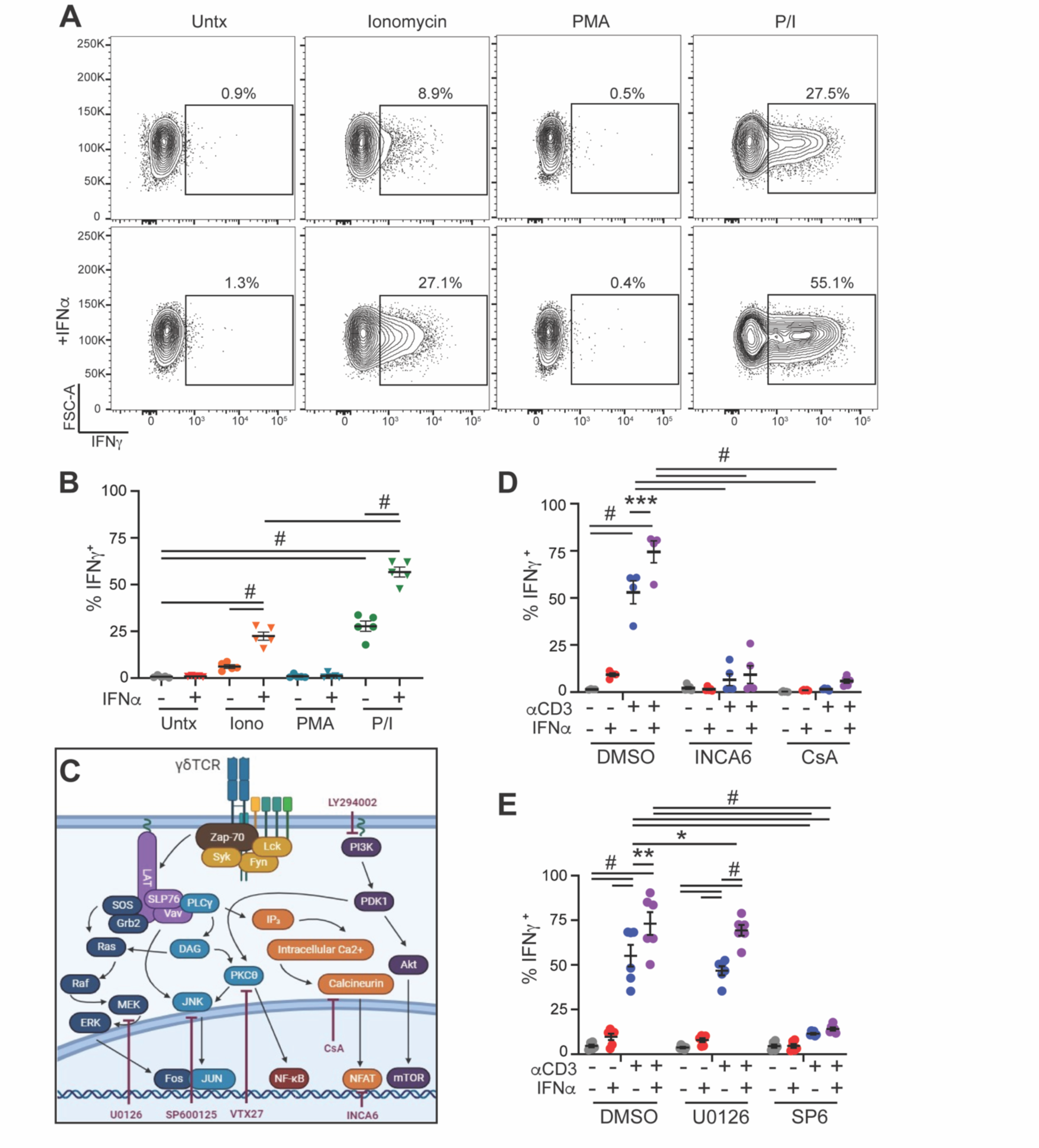
TCR activation of NFAT and JNK are required forγδ IEL IFNγ production. (A) Representative flow cytometry plots and (B) frequency of IFNγ^+^ freshly-isolated γδ lELs treated with 4 µg/mL ionomycin, 40 ng/mL PMA, or both (P/l) ± 10 ng/mL IFNα for 5 h. (C) Schematic of TCR signaling pathway including targets of pharmacological inhibitors. Percentage of IFNγ^+^ γδ lELs follow­ing 30 min pretreatment with DMSO, (D) 10 µM INCA6, 5 µM cyclosporin (CsA), (E) 10 µM SP600125 (SP6) or 10 µM U0126 for 30 min followed by 1 µg/mL αCD3, 10 ng/mL IFNα, or both for 5 h. Data are shown as mean ± SEM. Two independent experiments; n= 4-6 mice; *p<0.05, ***p<0.001, #p<0.0001; B: One-way ANOVA with Tukey’s multiple comparison test, D,E: Two-way ANOVA with Tukey’s multiple

To identify the specific downstream signaling components involved in γδ IEL IFNγ production, we next investigated the requirement for calcineurin/NFAT signaling using cyclosporin A (CsA) or INCA-6 (Figure 3C). Whereas CsA broadly inhibits calcineurin activity (39), INCA-6 more selectively prevents NFAT dephosphorylation by calcineurin (40). Use of either inhibitor completely abrogated cytokine production by γδ IELs following TCR activation, and as a result, IFNα was unable to elicit a response (Figure 3D, Supplementary Figure 3A).

Next, we sought to determine which pathways downstream of PMA contribute to IFNα-mediated IFNγ production. PMA activates ERK and JNK pathways; therefore, we treated γδ IELs with SP600125 and U0126, which selectively inhibit JNK1/2 and MEK1/2, respectively (41, 42).

Inhibition of JNK, but not MEK, prevented TCR-mediated cytokine production by γδ IELs (Figure 4E, Supplementary Figure 3B). Since CD28/B7 ligation induces PI3K signaling, yet CD28 is not expressed on γδ IELs (37, 43), the extent to which PI3K signaling contributes to γδ IEL effector function in response to a TCR agonist remains unclear. Pharmacological inhibition of PI3K reduced TCR-mediated cytokine production, yet this partial responsiveness to TCR agonist was sufficient to induce IFNα-induced IFNγ production (Supplementary Figure 3C,D). Although PKCθ can activate JNK in conventional T cells (44), inhibiting this pathway had no effect on γδ IEL IFNγ production.

**Figure 4:**
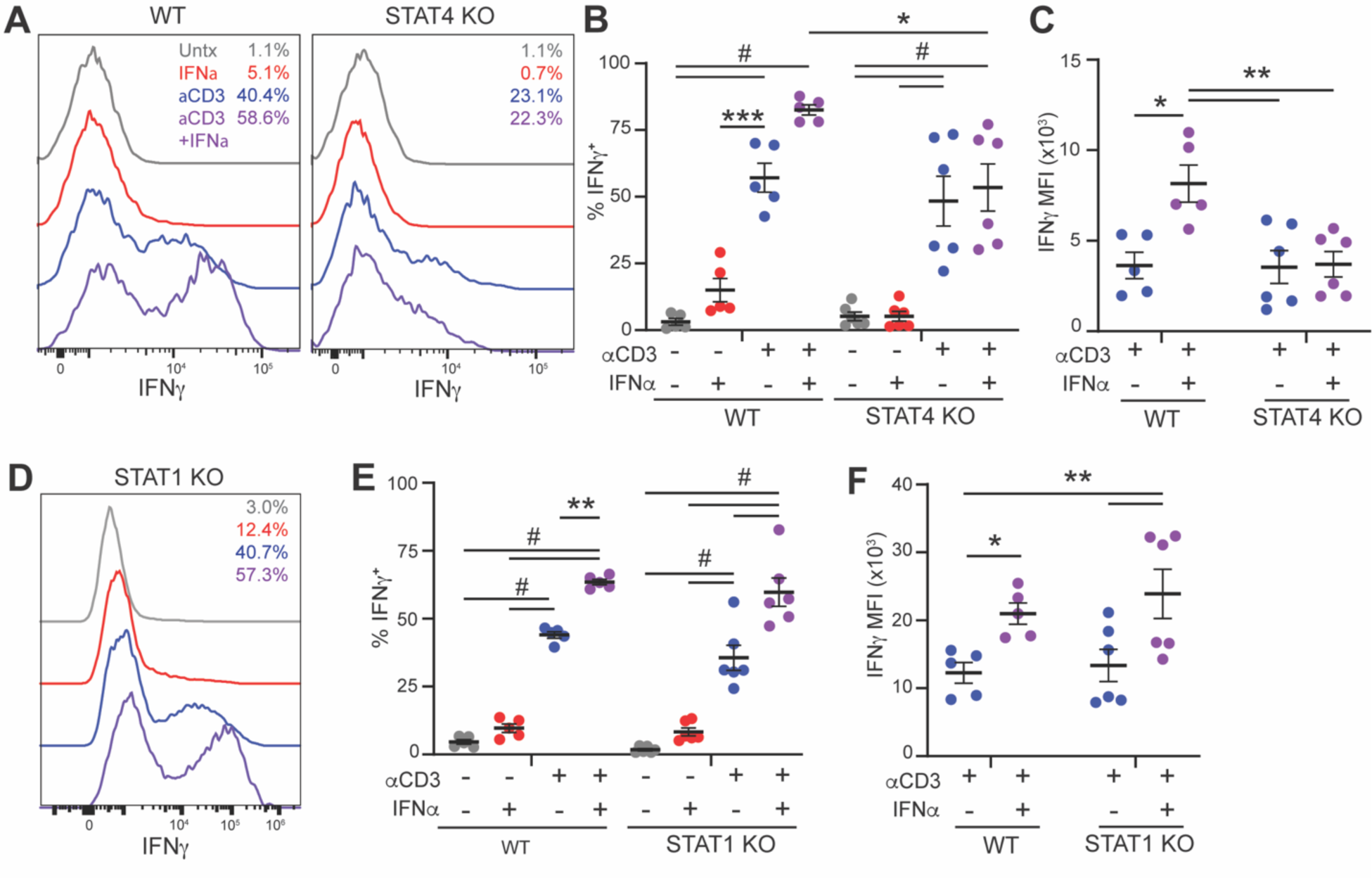
STAT4, but not STAT1, is required for enhanced. γδ **IEL IFNγ production in response to type I IFN.** (A,D) Representative histograms of IFNγ^+^ γδ lELs isolated from WT, STAT4 KO, or STAT1 KO mice, cultured ex *vivo,* and treated with 1µg/mL αCD3, 10 ng/mL IFNα, or both for 5 h. (B,E) Frequency or (C,F) MFI of IFNγ^+^ γδ lELs isolated from (B,C) STAT4 KO or (E,F) STAT1 KO mice compared to WT controls. Data are shown as mean ± SEM. Two independent experiments; n= 5-6 mice; *p <0.05, **p <0.01, #p <0.0001. B,C,E,F: Two-way ANOVA with Tukey’s multiple comparison test.

From these studies, we conclude that NFAT and JNK play a critical role in mediating cytokine production in response to TCRγδ activation. JNK phosphorylates c-Jun to allow binding to Fos and formation of the AP-1 complex, which cooperates with NFAT to modulate the transcriptional activation of *Ifng* promoter in response to TCR stimulation (45). Based on the known role of AP-1 complex in chromatin remodeling (46), we posit that the TCR signal creates permissive conditions to allow the binding of STAT proteins to the *Ifng* promoter, thus amplifying the expression of this cytokine in response to IFNα co-stimulation.

### STAT4 mediates enhanced γδ IEL IFNγ production in response to type I IFN

Activation of STAT1 and STAT4 are well-characterized in their regulation of type I IFN signaling in innate and adaptive host defense and IFNγ production (10, 23–25, 27). Therefore, we assessed the relative contribution of these two transcription factors in type I IFN-induced IFNγ production in γδ IELs. As expected, IFNα induced the phosphorylation of both STAT1 and STAT4 (Supplementary Figure 4A,B). We found that loss of STAT4 abrogated the co-stimulatory effect of IFNα in γδ IELs, whereas STAT1 was dispensable for IFNα-mediated IFNγ expression (Figure 4). These findings reflect previous reports demonstrating the requirement for STAT4 in type I IFN-induced IFNγ production in conventional CD8 T cells (10). Since IL-12 also promotes STAT4-mediated IFNγ in CD8 T cells (23), we asked whether IL-12 also enhances γδ IEL IFNγ production. We find that the frequency of IFNγ^+^ γδ IELs following IL-12 co-stimulation was similar to that observed in response to co-stimulation with IFNα (Supplementary Figure 4C). However, the amount of IFNγ produced by IL-12-treated γδ IELs in the presence of a TCR agonist was significantly higher than that following treatment with IFNα due to sustained STAT4 phosphorylation in response to IL-12 (Supplementary Figure 4D-F). In summary, we find that *ex vivo* cultured γδ IELs respond to type I IFN in a manner similar to conventional CD8 T cells in which TCR signaling synergizes with STAT4 to enhance IFNγ production.

### Activation of an IFNAR-mediated antimicrobial response in γδ IELs is regulated independently of tonic TCR signaling *in vivo*

Prior reports of constant TCRγδ triggering at steady-state (21, 22) coupled with our findings that a suboptimal TCR signal allows type I IFN induction of IFNγ *ex vivo* (Figure 2F) led us to ask whether tonic TCR signaling is sufficient to drive IFNα co-stimulation *in vivo*. We performed transcriptomic analysis of γδ IELs following acute, systemic IFNα administration of mice that were previously treated with control IgG or anti-TCRγδ (UC7) to induce TCR internalization.

PCA analysis showed that the transcriptome of γδ IELs segregated based on PBS or IFNα treatment, regardless of exposure to anti-TCRγδ (Figure 5A). We next identified differentially expressed genes among γδ IELs isolated from all conditions compared to IgG-PBS controls. Interestingly, inhibition of TCR signaling (UC7-PBS) minimally influenced gene expression, yet did lead to an upregulation of genes associated with NK receptor activity and cytotoxicity (*Klrc1*, *Klrc2*), suggesting thatÿgd IELs may upregulate NK receptors in the absence of a basal TCR signal to maintain immunosurveillance (Figure 5B,C).

**Figure 5:**
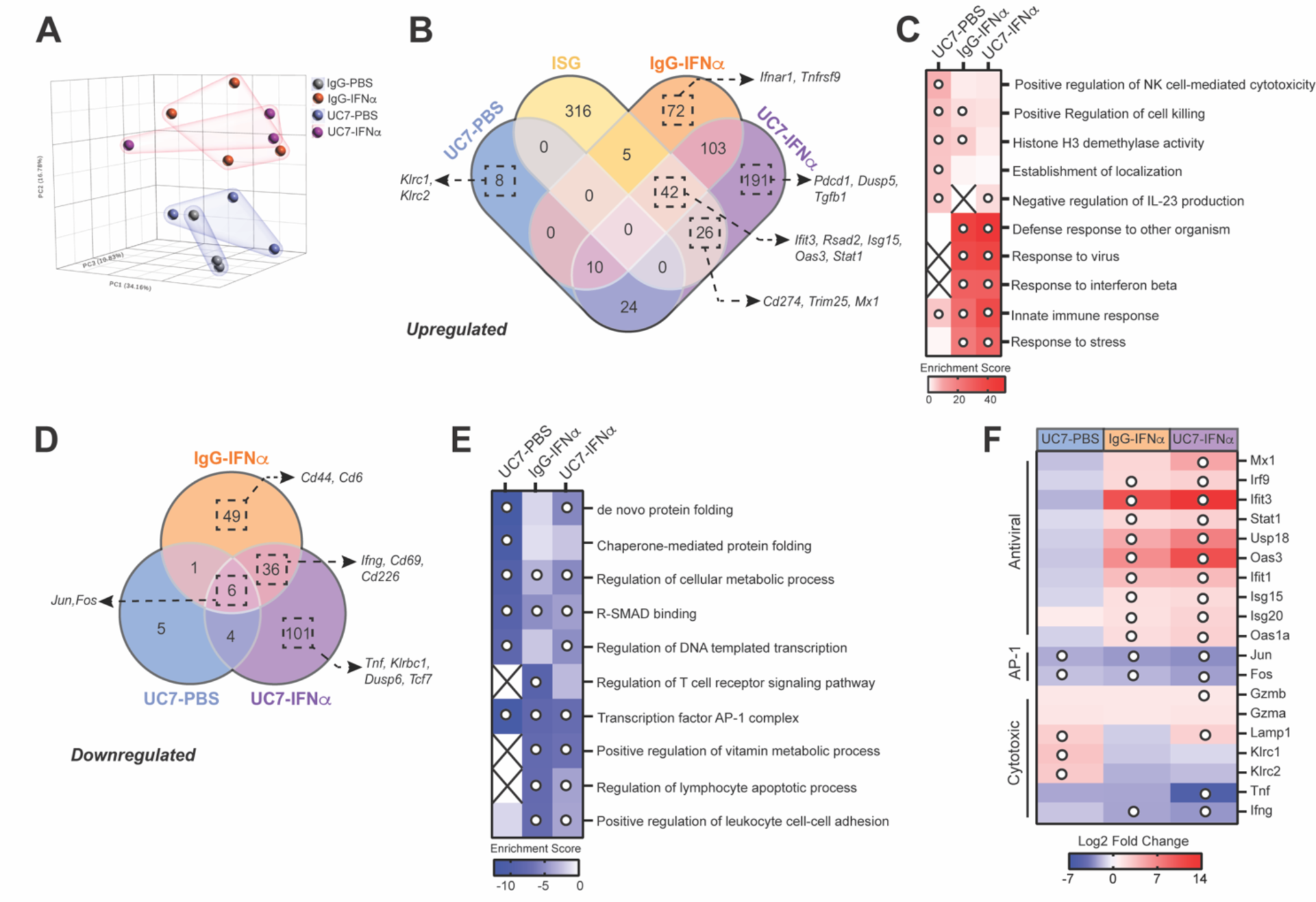
Type IIFN induces γδ IEL antiviral ISG expression in a TCR-independent manner. RNA sequencing of γδ lELs isolated from mice treated with IgG or anti-TCRγδ blocking antibody (UC7-13D5) 48 h prior to administration of PBS or 1 µg IFNα i.v. (A) Principal coordinate analysis of the γδ lELtranscriptome based on treatment conditions. (B) Upregulated genes among the various treatment conditions relative to IgG-PBS control were compared to ISGs upregulated in splenic γδ T cells following 2 h systemic IFNα treatment. The overlap among groups was summarized as a Venn diagram. Representative genes in each group indicated by arrows. (C) Gene set enrichment analysis of upregulated genes comparing between IgG-IFNα-, UC7-PBS-, or UC7-IFNα-treated groups and IgG-PBS control. “X” indicates conditions with no enrichment. (D) Venn diagram summarizing downregulated genes among the various treatment conditions relative to IgG-PBS control. Representative genes in each group indicated by arrows. (E) Gene set enrichment analysis of downregulated genes comparing between IgG-IFNα-, UC7-PBS-, or UC7-IFNα-treated groups and IgG-PBS control. “X” indicates conditions with no enrichment. (F) Heatmap indicating log-’2 fold change of selected genes involved in antiviral or cytotoxic effector response. n= 3-4 mice; C,E: White circle indicates p <0.05; F: White circle indicates FDR <0.05.

We then cross-referenced all upregulated genes against a group of ISGs previously shown to be rapidly upregulated in splenic γδ T cells following systemic IFNα treatment (34). 42 genes overlapped all three groups (IgG-IFNα, UC7-IFNα and γδ splenocyte ISGs), including those associated with a type I IFN-mediated antiviral signature (*Ifit3, Rsad2, Isg15, Oas3, Stat1*)(47, 48), demonstrating that activation of an IFN-induced innate immune program occurs in a TCRγδ-independent manner (Figure 5B,C).

To further dissect the contribution of tonic TCR signaling on the γδ IEL response to IFNα, we assessed gene expression unique to IgG-IFNα or UC7-IFNα treatment. Among the 77 DEGs upregulated following IFNα treatment alone (IgG-IFNα), we found that the expression of genes encoding IFNAR and the activating receptor 4-1BB (*Ifnar1, Tnfrsf9*)(23, 24, 49) was increased, whereas other genes associated with T cell activation (*Cd44, Cd6*)(50) were downregulated in this treatment group (Figure 5B-E). Inhibition of basal TCR activation in the presence of IFNα (UC7-IFNα) resulted in differential expression of 292 genes, indicating that TCRγδ plays a greater role in regulating the response to an exogenous stimulus than at steady-state (Figure 5B,F). Unexpectedly, genes that we propose to be critical for γδ IEL IFNγ production, specifically the AP-1 complex (*Jun, Fos*), were downregulated across all experimental conditions relative to control (Figure 5D-F). Consistent with this, *Ifng* expression was similarly reduced indicating that tonic TCRγδ activation is not sufficient to synergize with IFNAR to promote *Ifng* expression *in vivo*. Together with the observed reduction in *Tnf* expression in UC7-IFNα-treated mice (Figure 5F), we conclude that type I IFN alone, or in conjunction with basal TCRγδ signaling, fails to induce pro-inflammatory cytokine production in γδÿIELs.

Notably, an increase in select ISGs was observed only in UC7-IFNα-treated mice (*Mx1, Trim25*) as was *Tgfb1,* an immunoregulatory molecule (51)(Figure 5B,F). Similar to type I IFN alone, combined treatment led to differential regulation of various genes associated with inhibition of T cell activation; some were downregulated (*Klrb1c*, *Dusp6*), while others including *Pdcd1* (PD1), *Cd274* (PDL1), and *Dusp5* were upregulated (Figure 5B,D). Based on the variable expression of activating and inhibitory genes, it is difficult to definitively determine the contribution of tonic TCR signaling to the overall activation state of γδ IELs in response to type I IFN. Despite this, it is evident that acute exposure to IFNα drives innate immune responses in γδ IELs in a TCR-independent manner, thus reinforcing the notion that γδ IELs serve as sentinels of the epithelial barrier.

### γδ IELs exhibit reduced TCR responsiveness thus limiting pro-inflammatory cytokine production *in vivo*

As tonic TCRγδ triggering was unable to induce IFNγ production in response to IFNαÿ*in vivo*, we next investigated whether exogenous TCR stimulation could surmount the necessary activation threshold to allow for IFNα-mediated co-stimulation. IFNγ-YFP reporter mice were treated with TCR agonist in the presence or absence of IFNα to assess IFNγ production *in vivo*. We observed that the frequency of IFNγ-YFP^+^ cells was increased among induced CD4 or CD8αβ TCRαβ IELs relative to natural CD8αα IELs, which responded poorly to TCR activation (Figure 6A,B, Supplementary Figure 5A). Strikingly, IFNα failed to enhance IFNγ production in any IEL subset when treated either concurrent with TCR stimulation or for two hours prior to IEL isolation (Figure 6A,B, Supplementary Figure 5B). These data suggest that even in response to activation of innate immune signaling, IEL IFNγ expression is tightly regulated within the mucosal microenvironment.

**Figure 6:**
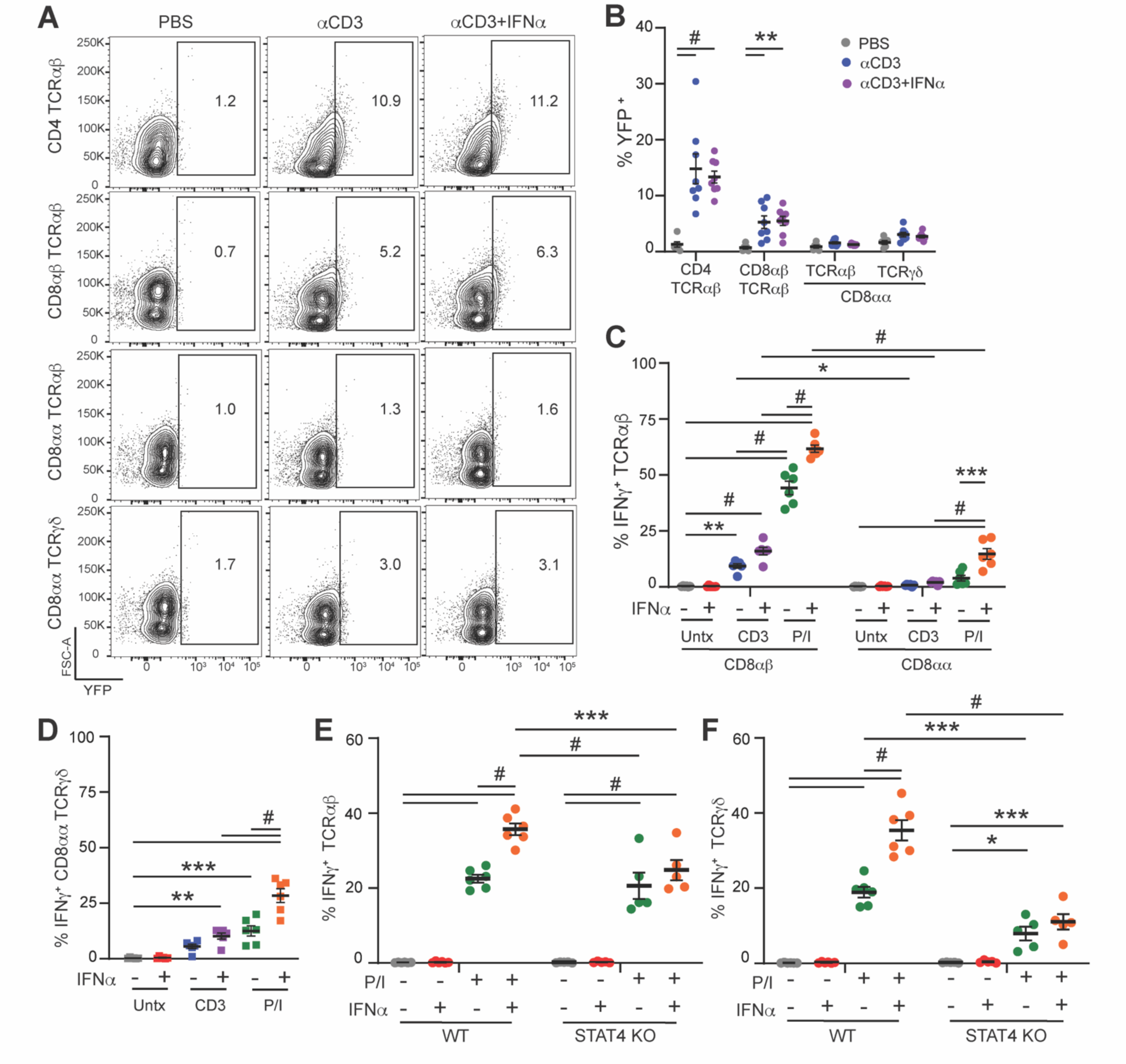
**Restricted TCR signaling in CDδαα lELs prevents IFNAR/STAT4-mediated IFNγ production *in vivo*** (A) Representative histogram showing (B) the frequency of IFNγ-YFP* CD4, CDδαβ, or CDδαα lELs in response to 24 h treatment with 25 µg αCD3 and/or 1 µg IFNα *in vivo.* To bypass proximal TCR signaling, freshly-isolated lELs were stimulated ex *vivo* with 1 µg/mL αCD3 or P/l ± 10 ng/mL IFNα for 5 h. The frequency of IFNγ+ CDδαβ or CDδαα lELs among (C) TCRαβ or (D) TCRγδ populations is shown. Freshly-isolated WT or STAT4-deficient lELs were treated with P/l ± of 10 ng/mL IFNα for 5 h and the frequency of IFNγ+ (E) TCRαβ or (F) TCRγδ lELs is shown. Data are graphed as mean ± SEM. Two independent experiments; n=5-6 mice; *p <0.05, ** p <0.01, ***p <0.001, #p <0.0001; B-E: Two-way ANOVA with Tukey’s multiple compari­son test.

The attenuated responsiveness of natural IELs to TCR stimulation has been reported by several groups. Interaction of the CD8αα homodimer with TL expressed on intestinal epithelial cells limits IEL effector response (52) and rewiring of IEL TCR signalosome dampens their response to activation (19). Based on the lack of IFNα co-stimulation in IELs *in vivo*, we asked whether bypassing proximal TCR signaling by activating freshly-isolated IELs with P/I differentially affects IFNγ production among the various IEL subsets. Activated, induced CD8αβÿTCRαβ IELs exhibited a more robust IFNγ response relative to natural CD8αα TCRαβ IELs, yet activation with P/I restored IFNα-induced IFNγ production in CD8αα TCRαβ IELs (Figure 6C). Similar findings were observed in CD8αα γδ IELs, in which the frequency of IFNγ^+^ cells was increased following treatment with IFNα in the presence of either a TCR agonist or P/I (Figure 6D). Whereas anti-CD3 alone did not induce significant IFNγ production in freshly-isolated CD8αα γδ IELs, the slight increase in IFNγ following IFNα co-stimulation suggests that, to a certain extent, the TCR activation threshold necessary to elicit IFNγ production may have been met.

Our data indicate that removing γδ IELs from the intestinal microenvironment and subjecting these cells to *ex vivo* culture likely provides a more permissive context for TCR responsiveness, and subsequently, increases the capacity to produce IFNγ in response to type I IFN. To determine if the requirement for IFNAR/STAT4-mediated IFNγ production is conserved among natural CD8αα IELs *in vivo*, we stimulated WT or STAT4-deficient CD8αα IELs with P/I and/or IFNα immediately after isolation. Consistent with our *ex vivo* culture data (Figure 4A- C), IFNα-mediated co-stimulation is abrogated in the absence of STAT4 in both CD8αα TCRαβ and CD8αα γδ IELs (Figure 6E,F). Interestingly, the frequency of IFNγ^+^ STAT4-deficient γδ IELs was reduced following P/I treatment compared to WT, suggesting that STAT4 may also contribute to IFNα-independent mechanisms of γδ IEL activation (Figure 6F). Collectively, these data demonstrate that natural IELs retain the capacity for enhanced IFNγ production through IFNAR/STAT4 signaling *in vivo*; however, reduced TCR responsiveness limits their ability to produce IFNγ. This may reflect an important regulatory mechanism to fine-tune mucosal immune responses while also preventing excessive IFNγ production within the epithelial compartment that could negatively impact barrier function (53, 54).

## Discussion

Despite exhibiting functional similarities between NK cells, we find TCR activation is required for IFNα-mediated IFNγ production in *ex vivo* cultured γδ IELs. Further investigation into the underlying molecular mechanisms of co-stimulation highlighted similarities between type I IFN activation of γδ IELs and conventional CD8 T cells. The TCR-dependent response to type I IFN *ex vivo* led us to investigate the components of the TCRγδ signaling pathway involved in driving IFNγ production. The use of various pharmacological inhibitors showed that γδ IELs are activated in a manner similar to conventional CD8 T cells, with NFAT and AP-1 driving pro-inflammatory gene expression (55). NFAT is necessary to induce rapid IFNγ production by CD8 T cells (55–57); however, uncoupling NFAT and AP-1 induces cytolytic effector cells to adopt an exhausted phenotype, indicative of reduced cytotoxic effector function (58). Since AP-1 signaling is critical, and NFAT alone is insufficient to induce IFNγ production, it is no surprise that inhibition of either NFAT or JNK signaling abrogates γδ IEL IFNγ production following activation *ex vivo*. Further, TCR-mediated γδ IEL pro-inflammatory cytokine production was blunted in the presence of PI3K inhibition, suggesting that PI3K signaling may contribute to activation of the AP-1 complex through JNK (59). These data demonstrate that TCR activation of *ex vivo* cultured γδ IELs induces IFNγ production through conserved intracellular signaling pathways akin to conventional CD8^+^ T cells, despite the lack of a clear co-stimulatory signal.

IFNAR/STAT signaling regulates the timing of innate and adaptive immune responses to viral infection, in which basal STAT4 expression promotes IFNγ production by NK cells early in infection, and STAT1 constrains CD8 T cells until antigen-specific effector T cells have been mobilized (23–25). Following antigen recognition, the upregulation of STAT4 facilitates effector CD8 T cell expansion and IFNγ production in response to IFNα/IL-12 co-stimulation. γδ IELs bridge innate and adaptive immunity with properties of both NK cells and conventional CD8 T cells. Unlike NK cells, exposure to type I IFN or IL-12 alone was incapable of driving IFNγ, yet the addition of a suboptimal TCR signal synergized with IFNα to enhance γδ IEL IFNγ production. We find that IFNAR signaling through STAT4 is required for γδ IEL IFNγ production, and despite reduced TCR responsiveness *in vivo*, bypassing proximal TCR signaling events confirmed that these pathways remain intact *in situ*. Taken together, we propose two potential mechanisms by which type I IFN amplifies IFNγ production in γδ IELs: (1) cooperative signaling between TCR- and IFNAR-induced transcription factors or (2) TCR stimulation increases chromatin accessibility to allow STAT4 binding to the IFNγ promoter.

Previous work has shown that NFAT and STAT4 both bind the promoter region of *Pdcd1* and induce active enhancer modifications that synergize to drive gene transcription (60), yet there is little evidence supporting cooperative binding of these transcription factors to promote *Ifng* expression. Moreover, the ability of IFNα to increase the frequency of IFNγ^+^ γδ IELs following ionomycin stimulation suggests that AP-1 activation is not required for STAT4 binding to the *Ifng* promoter. NK cells can rapidly produce IFNγ based on the constitutive expression of transcription factors that bind to *Ifng* regulatory elements located within accessible chromatin regions (61–63), whereas demethylation of a highly methylated *Ifng* locus correlates with gene transcription in T cells (64–66). We posit that under permissive conditions, TCR stimulation increases accessibility of the *Ifng* locus to enable STAT4-mediated IFNγ production in γδ IELs in response to IFNα. However, in the absence of a TCR signal *ex vivo*, or reduced TCR responsiveness *in vivo*, the *Ifng* locus remains inaccessible resulting in minimal γδ IEL IFNγ production.

Relatively little is known about the epigenetic landscape of IELs and how tonic, tissue-derived signals may influence chromatin accessibility. In invariant NK T cells (iNKT), a strong TCR signal leads to IFNγ production yet a weak signal induces histone acetylation at the *Ifng* locus without inducing gene transcription, suggesting that the strength of the TCR signal imprints future responsiveness to external stimuli (67). Whereas increased basal calcium flux among freshly-isolated γδ IELs provided the first indication of constant TCR triggering *in vivo* (21), more recent analyses reveal that only 6% of γδ IELs receive a TCR signal within 4 hours prior to isolation (22). As a result, the biological contribution of tonic TCRγδ triggering remains unclear. We observed limited differential gene expression among γδ IELs isolated from mice administered TCR blocking antibody; however, the upregulation of genes involved in regulating innate-like NK responses provides a potential alternative mechanism to maintain immune surveillance in the absence of TCR surface expression.

Many of the genes upregulated in response to IFNα contribute to host innate immune responses independent of TCR signaling; this is consistent with other known roles for γδ IELs in epithelial surveillance and antimicrobial function, many of which do not require TCR signaling (4, 14, 68, 69). Inhibition of TCR signaling further increased IFNα-induced antiviral gene transcripts in γδ IELs, suggesting that gene loci involved in innate immunity are either located in more accessible regions under homeostatic conditions or tonic TCRγδ signaling negatively regulates the response to type I IFN. We report that the inhibition of basal TCR signaling decreased *Ifng* transcripts in γδ IELs following IFNα exposure, leading us to ask, does tonic TCR signaling influence the epigenetic landscape of γδ IELs? It is possible that low level calcium signaling downstream of basal TCRγδ activation may prime the epigenetic modification of genes involved in a pro-inflammatory, adaptive-like response. However, the inability of γδ IELs to induce pro-inflammatory cytokine production in response to TCR agonist or type I IFN reflects a tightly controlled effector response. Future studies focusing on the epigenetic regulation of IEL function may provide critical insight regarding the propensity of, and conditions under which, these sentinel lymphocytes mount an innate-like versus an adaptive-like effector response.

In contrast to induced CD8αβ IELs, we find that natural CD8αα IELs remain unable to produce IFNγ even after prolonged exposure to TCR agonist *in vivo*. Recent work investigating the TCR signalosome of natural IELs shows that many signaling components are downregulated or substituted with less active proteins compared to conventional T cells (19). Taking this into account, previous work by Swamy et al., showed that *in vivo* administration of a TCR agonist drove IEL production of type I, II and III interferon which was then able to confer antiviral protection against murine norovirus (MNV)(14). However, infection with MNV.CR6, a potent inducer of type I IFN, did not affect γδ IEL number or cytokine production (70). Similar this report, we were unable to detect IFNγ protein expression in freshly-isolated γδ IELs following anti-CD3 treatment by intracellular staining or using IFNγ-YFP reporter mice. Based on the attenuation of responsiveness to TCR stimulation within the natural IEL compartment, we posit that virus-specific, induced CD8αβ IELs or lamina propria T cells may function as the primary responders to TCR agonist (71, 72) and contribute to the production of type I IFN that can subsequently promote an innate, antiviral program in γδ IELs as part of a bystander response (73).

Thus, there are several outstanding questions relating to how γδ IEL effector function is regulated at steady-state or in response to disease. Our study highlights some of the limitations involved in addressing these questions based on (1) the attenuated of responsiveness to TCR stimulation *in vivo* and (2) *ex vivo* culture models do not necessarily faithfully represent the activation state of γδ IELs *in vivo*. We hypothesize that γδ IEL activation in response to cytokine is tightly regulated by a combination of epithelial-derived signals and cytokine receptor signaling. Further investigation is needed to elucidate the capacity for TCR responsiveness among γδ IELs *in vivo* and the biological context that allows these cells to exceed their activation threshold, whether this occurs through immunologic, metabolic or epigenetic regulation.

Additional studies regarding the underlying mechanisms involved in regulating γδ IEL responsiveness may be helpful in developing strategies to modulate conventional T cell activation and limit aberrant inflammation in the context of autoimmunity.

## Acknowledgments

The authors would like to thank Sergei Kotenko and Tessa Bergsbaken for providing the STAT1 and STAT4 KO mice in addition to their thoughtful suggestions on the manuscript. Cell sorting was performed at the NJMS Flow Cytometry and Immunology Core Laboratory and supported by National Institute for Research Resources Grant S10RR027022. Schematics were designed using Biorender.

## Author Contributions

M.F. and L.J. designed and performed experiments and analyzed the data. M.F. wrote the manuscript. K.L.E. conceived the study, supervised the research and wrote the manuscript. All authors approved the final manuscript.

## Abbreviations

DAG: diacylglycerol
DEG: differentially expressed gene
IEL: intraepithelial lymphocyte
IFNAR: IFNα/β receptor
iNKT: invariant NK T cell
IRF: IFN regulatory factor
ISG: IFN stimulated gene
MNV: murine norovirus
WT: wildtype

---

**Supplementary Figure 1:**
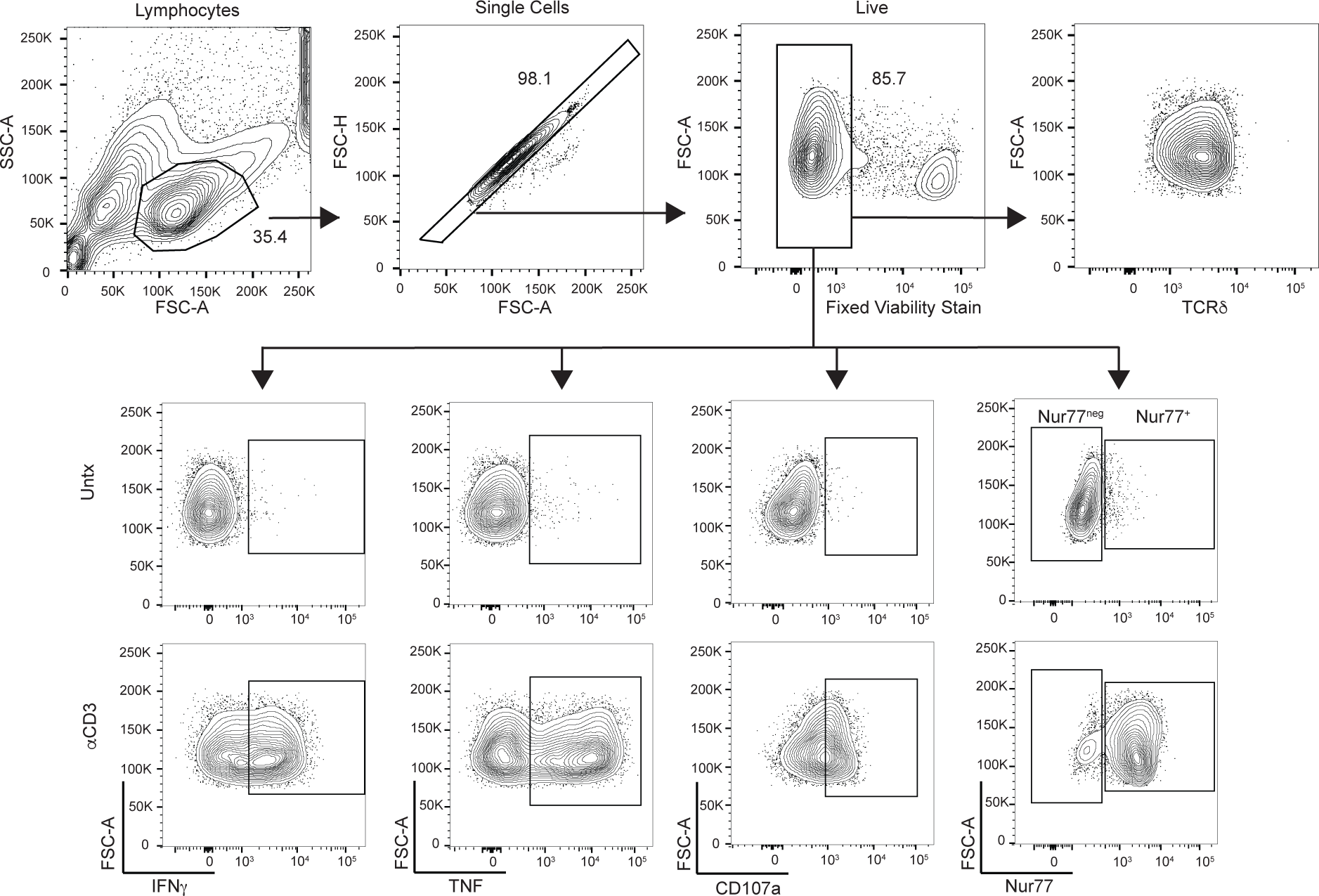
Gating strategy for *ex vivo* cultured. γδ **IELs.** Sorted γδ IELs were sequentially gated based on size, singlets, and viability. TCRδ expression within the cultured population is shown in addition to the representative gating strategy for IFNγ^+^, TNF^+^, CD107a^+^, or Nur77^+^ γδ IELs in the presence or absence of αCD3 stimulation (1μg/mL, 5 h).

**Supplementary Figure 2:**
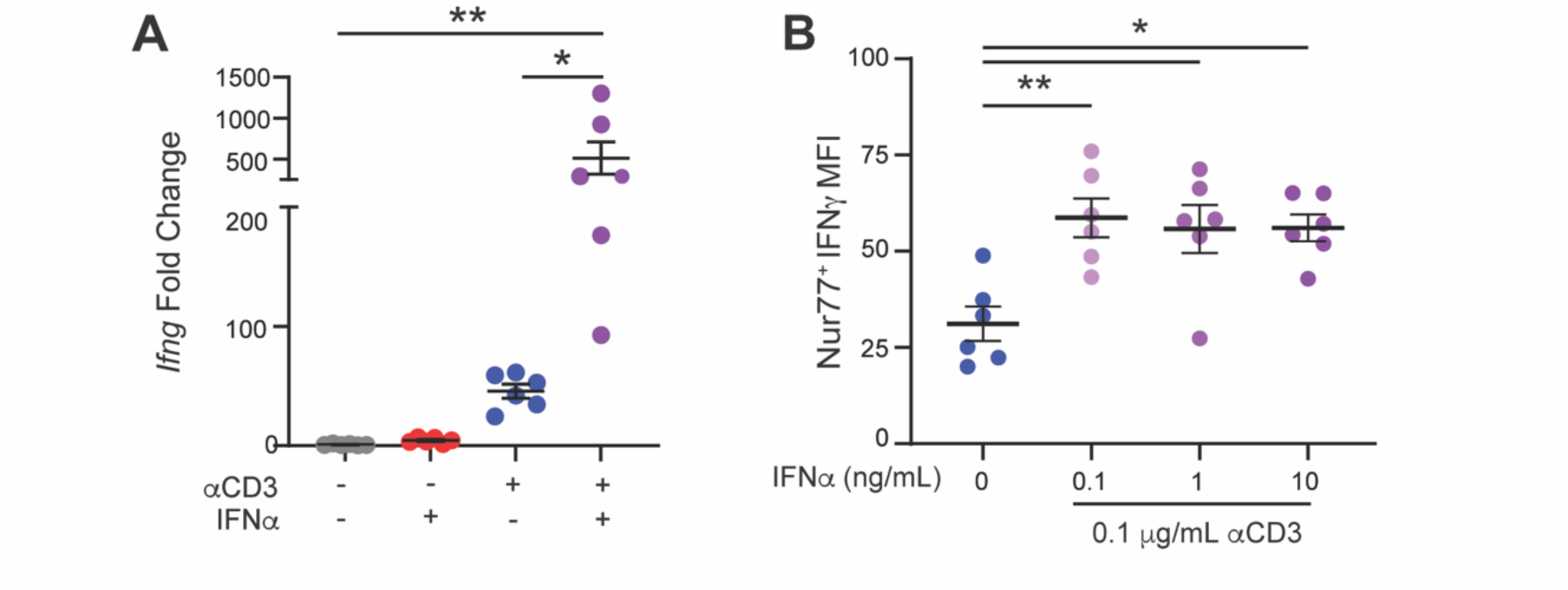
Type IIFN co-stimulation increases the transcription of *Ifng* in *ex vivo* cultured. γδ **lELs and promotes** IFNγ **production at low concentrations.** (A) Quantitative PCR was performed on RNA isolated from *ex vivo* cultured γδ lELs treated with 1µg/mL αCD3, 10 ng/mL IFNα, or both for 5 h. ΔΔCT was normalized to *Gapdh* and fold change compared to untreated samples is shown. (B) *Ex vivo* cultured γδ lELs were stimulated with 0-100 ng/mL IFNα and 0.1 µg/mL αCD3 for 5 h and the MFI of IFNγ among Nur77^+^ γδ lELs is shown. Data are graphed as mean ± SEM. Data are shown as mean ± SEM. Two independent experiments; n=5-6 mice. *p <0.05, **p <0.01. One-way ANOVA with Tukey’s multiple comparison test.

**Supplementary Figure 3:**
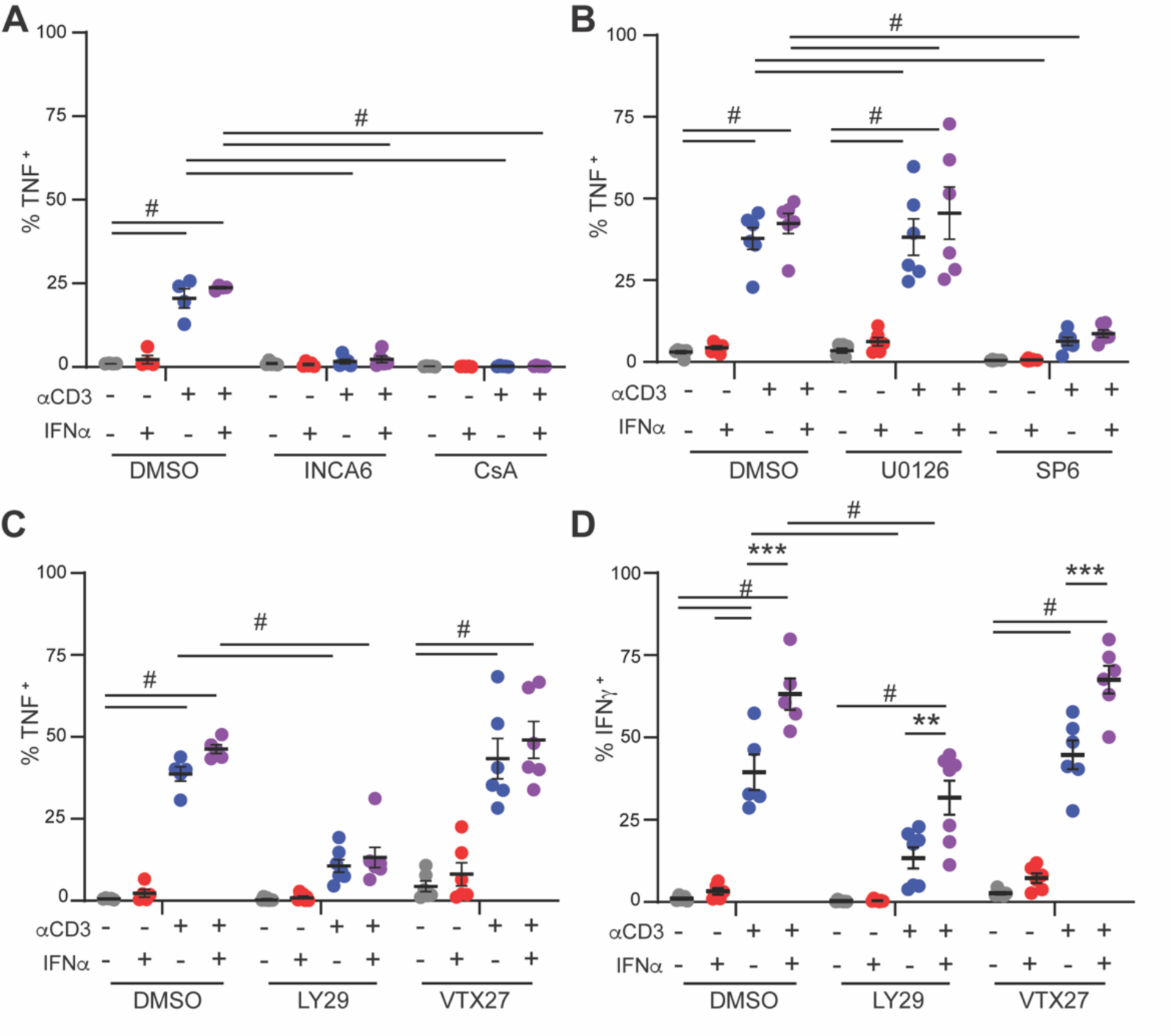
Signaling through PI3K or PKCΘ is not essential for IFNα- mediated IFNγ production in. γδ **lELs.** *Ex vivo* cultured γδ lELs were pretreated with DMSO, 10 µM INCA6, 5 µM cyclosporin (CsA), 10 µM SP600125 (SP6), 10 µM U0126, 10 µM LY294002 (LY29), or 1 µM VTX27 for 30 min followed by 1 µg/mL αCD3, 10 ng/mL IFNα, or both for 5 h and the frequency of (A-C) TNF* or (D) IFNγ* γδ lELs is shown. Data are represented as mean ± SEM. Two independent experiments; n= 5-6 mice; **p <0.01, ***p <0.001, #p<0.0001; Two-way ANOVA with Tukey’s multiple comparison test.

**Supplementary Figure 4:**
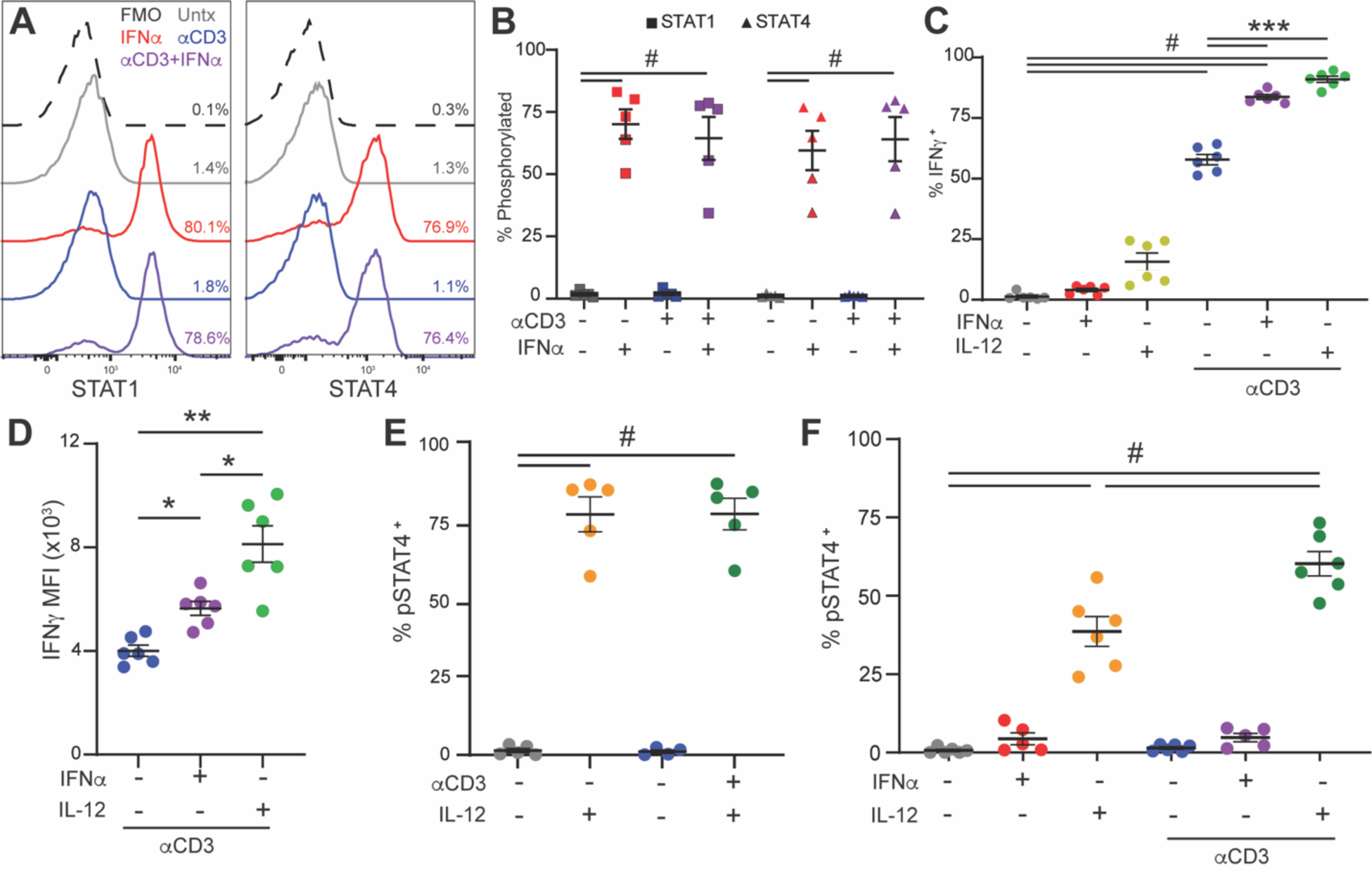
IFNAR and IL-12 both function as co-stimulatory signals to enhance IFNγ production in. γδ **lELs.** Representative histograms of (A) STAT1 or (B) STAT4 phosphorylation in ex *vivo* cultured γδ lELs following treatment with 1 µg/mL αCD3, 10 ng/mL IFNα, or both for 30 min. (C) Frequency of phospho-STAP γδ lELs. (D) Frequency of IFNγ^+^ or (E) IFNγ MFI of γδ lELs treated with 10 ng/mL IFNα or 10 ng/mL IL-12 ± 1µg/mL αCD3 for 5 h. Frequency of phospho-STAT4^ł^ γδ lELs after IL-12 treatment for (E) 30 min or (F) 5 h. Data are represented as mean ± SEM. Two independent exper­iments; n=5-6 mice; *p <0.05, **p <0.01, ***p <0.001, #p <0.0001; B: Two-way ANOVA with Tukey’s multiple comparison test. C-F: One-way ANOVA with Tukey’s multiple comparison test.

**Supplementary Figure 5:**
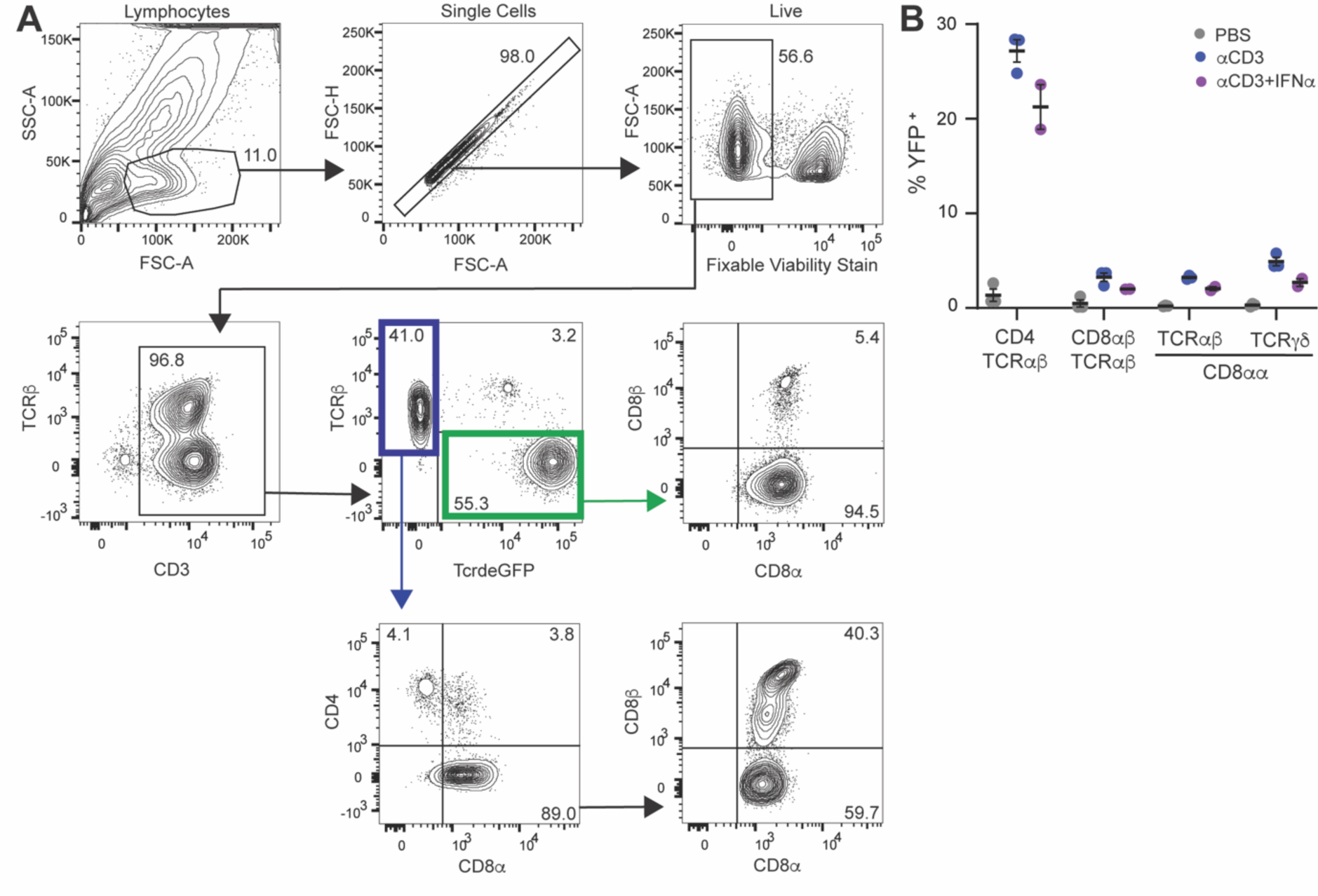
Acute IFNα exposure in conjunction with TCR stimulation does not induce IEL IFNγ production. *in vivo.* (A) lELs were gated based on size, singlets, and viability. CD3^ł^ cells were subsequently gated on TCRαβ (blue) or TCRγδ (green) expression after which TCRαβ lELs were further separated by CD4, CD8α, or CD8β expression. TCRγδ lELs were gated on CD8αα. (B) Frequency of IFNγ-YFP* CD4, CD8αα, or CD8αβ lELs in response to 24 h treatment with 25 µg αCD3 with or without an acute 2 h exposure to 1 µg IFNα *in vivo.* Data are representative of one experiment, n=2-3mice. Mean ± SEM is shown.

